# AND-gate fluorescence prodrug of the self-immolative delivery for overcoming multidrug resistance

**DOI:** 10.1101/2025.06.16.660040

**Authors:** Jun Liu, Zhengqin Liu, Yifen Yang, Yanli Long

## Abstract

The principle goal of clinical tumor treatment is to improve the therapeutic outcomes in the clinic while suppressing undesired purgatorial side-effects of anticancer drugs. However, its clinical usage of fluorescence prodrug was significantly hampered with some critical issues such as water-solubility, blood circulation, therapeutic efficacy, inevitable leakage, and side effects simultaneously. Herein, based on the strategy of the unique grafting structure, we report H_2_O_2_ activatable phototheranostic probes targeting the Golgi in drug-induced kidney injury tissues of mice with hypertension. The drug release study in vitro demonstrated that LGM-XL can diagnose cancer effectively by monitoring fluctuations of H_2_O_2_ in vivo through fluorescence imaging. More importantly, the prodrug LGM-XL could detect and image endogenous H_2_O_2_ in inflamed mice in situ, while effectively avoiding drug leakage in blood. The novel multifunctional drug delivery system enhanced not only Golgi oxidative stress in the pathophysiology of hypertension by stimulation of H_2_O_2_ but also the molecular mechanism for the diagnosis of inflamed-related diseases (e.g., drug-induced liver injury and the liver repair).

## 1. Introduction

Hypertension is a cardiovascular disease (CVD), contributing to an elevated incidence of endothelial dysfunction events by high blood pressure.^[13, 23]^ It is associated with the combined effects of genetic and environmental factors in this high-risk group of patients.^[3]^ Hypertension is the main cause of the high mortality and the threatening complications for public health in the world.^[4]^ As a multifactorial disorder, accumulating evidence suggests that oxidative stress play a crucial role in the pathological process of hypertension, which is an imbalance between antioxidant capacity and the production of reactive oxygen species (ROS) within the vasculature. ^[53, 63]^ In addition, Golgi oxidative stress could contribute to the vascular pathology and other organ damage associated with hypertension by expressed homodimeric enzyme. ^[7]^ However, thepotential therapeutic targets and the pathological process of essential hypertension remain unclear in the Golgi apparatus. Therefore, it is imperative to explore new the molecular mechanism and function of Golgi oxidative stress in the pathophysiology of hypertension.

Since the Golgi apparatus (GA) acts as a trafficking hub for packing, transport and sorting of proteins synthesized in the endoplasmic reticulum (ER).^[83, 93]^ Golgi apparatus is a pivotal organelle in most eukaryotic cells, which would be completely disassembled during the process of Golgi oxidative stress response under the situation of insufficient capacity.^[10-12]^ In addition, Golgi stress is observed in a wide range of kinds of diseases along with extensive ROS production, especially hydrogen peroxide (H_2_O_2_), an indicator of oxidative stress.^[13-15]^ The abnormal H_2_O_2_ level was acted as a signaling molecule and produced by resident proteins in physiological and pathophysiological processes of Golgi apparatus.^[16-17]^ Moreover, accumulating evidence suggests that Golgi-located H_2_O_2_ plays a central role in the pathogenesis of hypertension by inducing endothelial dysfunction.^[18-20]^ Therefore, complete understanding of the detailed mechanisms of Golgi stress response could reveal the direct relevance between hypertension and H_2_O_2_ levels in situ in real time.

Stimuli-responsive fluorogenic prodrugs can be engineered by using different imaging techniques.^[21-25]^ However, The success of prodrug was significantly hampered with the low stability, host toxicity, poor drug loading and short retention in the targeted tumor tissues. Moreover, the drug-induced liver injury (DILI) remains a challenge in drug discovery and clinical use, which is closely associated with the oxidative stress. Golgi oxidative stress could contribute to the vascular pathology and other organ damage associated with hypertension by expressed homodimeric enzyme. ^[26-28]^ Meanwhile, it is hypothesized that selective Golgi oxidative stress could be a strategy to provide cognitive benefits to cardiovascular disease (CVD) patients while suppressing undesired purgatorial side-effects of anticancer drugs and prolonging circulation in blood vessels. Thus, novel therapies are urgently needed to overcome these obstacles regarding the development of fluorogenic prodrugs.

Based on the above backgrounds, we are committed to develop a multifunctional theranostic probe (LGM-XL), which was designed and constructed by introducing 4-hydroxy group moiety of Golgi-targeted naphthalimide fluorophore with monoboronate as a well-known H_2_O_2_ reaction site, 5’-deoxy-5-fluorouridine (5′-DFUR) as anti-tumor drug and 4-dimethylaminoazobenzene-4′-carboxylic acid (DABCYL) as the quencher,^[29]^ by enhancing sensitivity and bioavailability for improved drug delivery. LGM-XL as a turn-on mode with ultra-high specificity and fast responses for Golgi H_2_O_2_ in cancer cells during oxidative stress. We demonstrated the information on the generation and distribution of H_2_O_2_ activity in the Golgi apparatus. Moreover, based on the lower cytotoxicity, probe LGM-XL was successfully applied to examine the Golgi H_2_O_2_ level in mice with hypertension in this process. Utilizing this probe, we revealed the mechanism of drug-induced liver injury and the liver repair processes. And most importantly, the framework could overcome the stability in the blood circulation and achieving drug release in acidic tumor environments. As such, this work favoringly disclosed the probe was a valuable new tool for visualizing Golgi H_2_O_2_ fluctuation and studying diagnosis method, which has not been investigated to date.

**Scheme 1.**
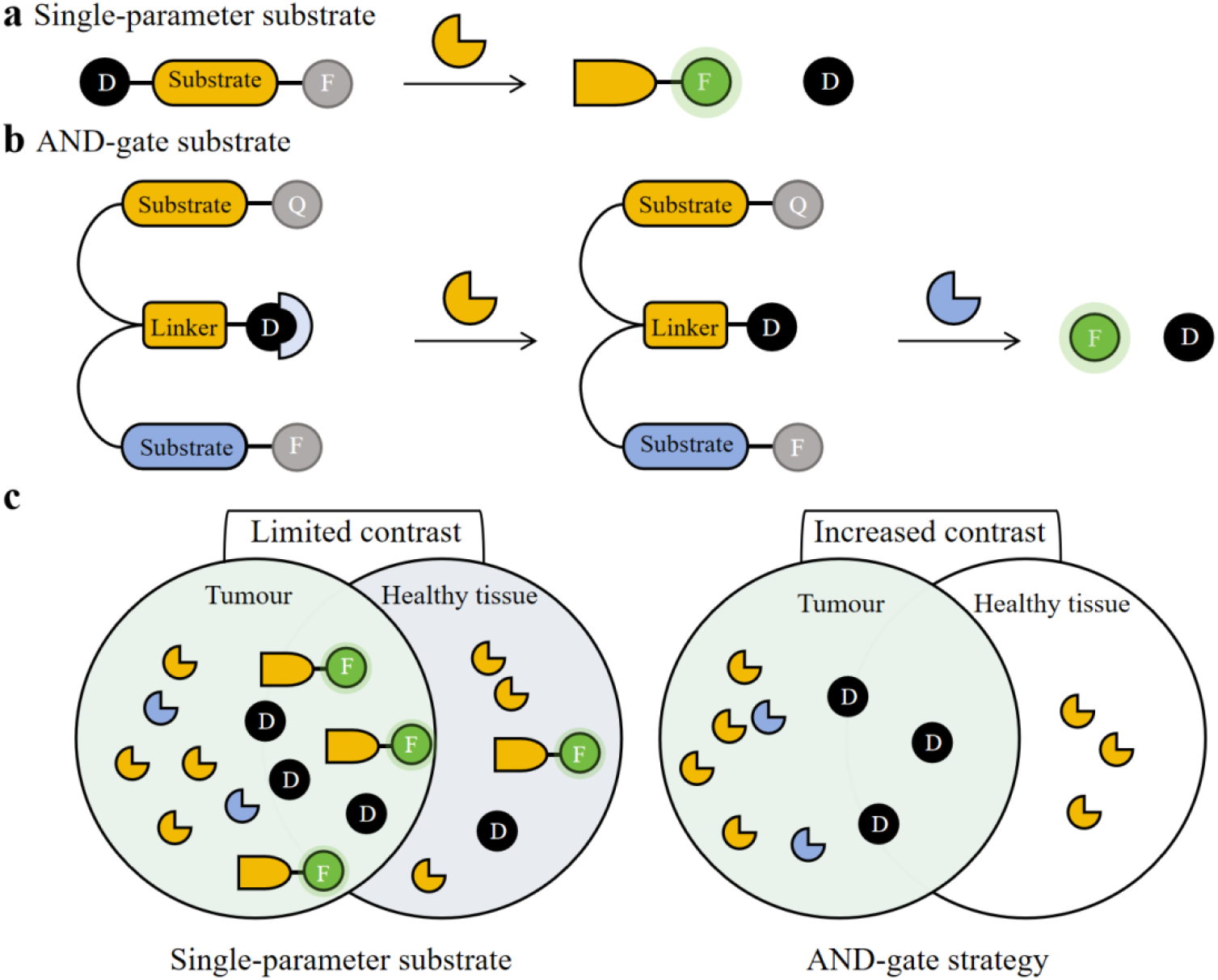
Schematic representation of the reaction of probe with Golgi H_2_O_2_. Q, quencher; F, fluorophore; D, drug. Comparison of the single-parameter and AND-gate probes in tumour and normal tissues.

## 2. Experimental section

### 2.1 Materials and Instruments

Solvents and chemicals were purchased from major commercial suppliers and applied directly in the experiment without further purification unless otherwise noted. Silica gel (Silicycle, 300−400 mesh) was prepared for column chromatography. Inorganic salts (MgCl_2_, CaCl_2_, KCl, NaCl, ZnCl_2_), reactive oxygen species (H_2_O_2_, TBHP, HClO, KO_2_), biothiols (cysteine), nitrogen oxide (NO), and cysteine (Cys) were prepared in aqueous solutions. All NMR spectra for the target compounds were recorded on a Bruker Advance-500 spectrometer with chemical shifts reported as ppm (in DMSO-d6, tetramethylsilane as the internal reference). Electrospray ionization mass spectroscopy (ESI-MS) data were measured with Bruker Esquire 3000 plus mass spectrometer. The fluorescence spectra were performed on a F-7000 fluorescence spectrophotometer (Hitachi). Living cell imagings were investigated through an Olympus FV1000-MPE multi-photon laser scanning confocal microscope (Japan). All the optical measurements were performed at room temperature. The synthetic route toward probe (LGM-XL) and analytical data were provided in supporting information and the molecular probes of interest for this study was outlined in Scheme 1.

### 2.2 Cell culture

Hela cells were cultured in DMEM medium (high glucose) supplemented with 10% FBS and 1% penicillin/ streptomycin (penicillin: 10,000 U.mL^−1^, streptomycin: 10,000 U.mL^−1^) under 5% CO_2_ environment stored for 12 h in a humidified atmosphere at 37 °C. Before imaging, the growth medium was removed and the cells were washed with PBS buffer (0.5 mL) for three times. Cells were passaged at about 80 % cell confluency using a 0.25% trypsin solution. Confocal fluorescence imaging was performed on confocal microscope (Leica SP8, Germany) using 488 nm excitation and the fluorescence emission windows were set as 520–560 nm.

### 2.3. Hypertension model mice

All animal studies were performed in agreement with the National Institutes of Health on the use of experimental animals (China), and were approved by the Animal Ethics Committee of Zunyi Medical University. 8-Weekold female BALB/c mice were supplied by Western Biomedical technology Lab Animal Co., Ltd. (Chongqing, China). In order to avoid the interference on imaging, the animals were kept in a controlled environment with consistent temperature (22 ± 2 °C) and humidity (50 ±10%) under the whole imaging process.

## 3. Results and Discussion

### 3.1 Screening and optimization of probe LGM-XL

We initially assessed the responsive ability of probe toward H_2_O_2_ by fluorescence spectrum in PBS buffer. As illustrated in Figure 1a, in the absence of H_2_O_2_, the fluorescence of the solution hardly changes under excitation at 480 nm. After the addition of H_2_O_2_ (160 μM), the fluorescence signal of probe showed a remarkable enhancement trend, indicating the phenylboronic group recognition group exhibited high reactivity for H_2_O_2_. Subsequently, it could be seen from Figure 1b that the fluorescence signal of probe LGM-XL at approximately 535 nm was lighten-up after the addition of different concentrations of H_2_O_2_.

**Figure 1.**
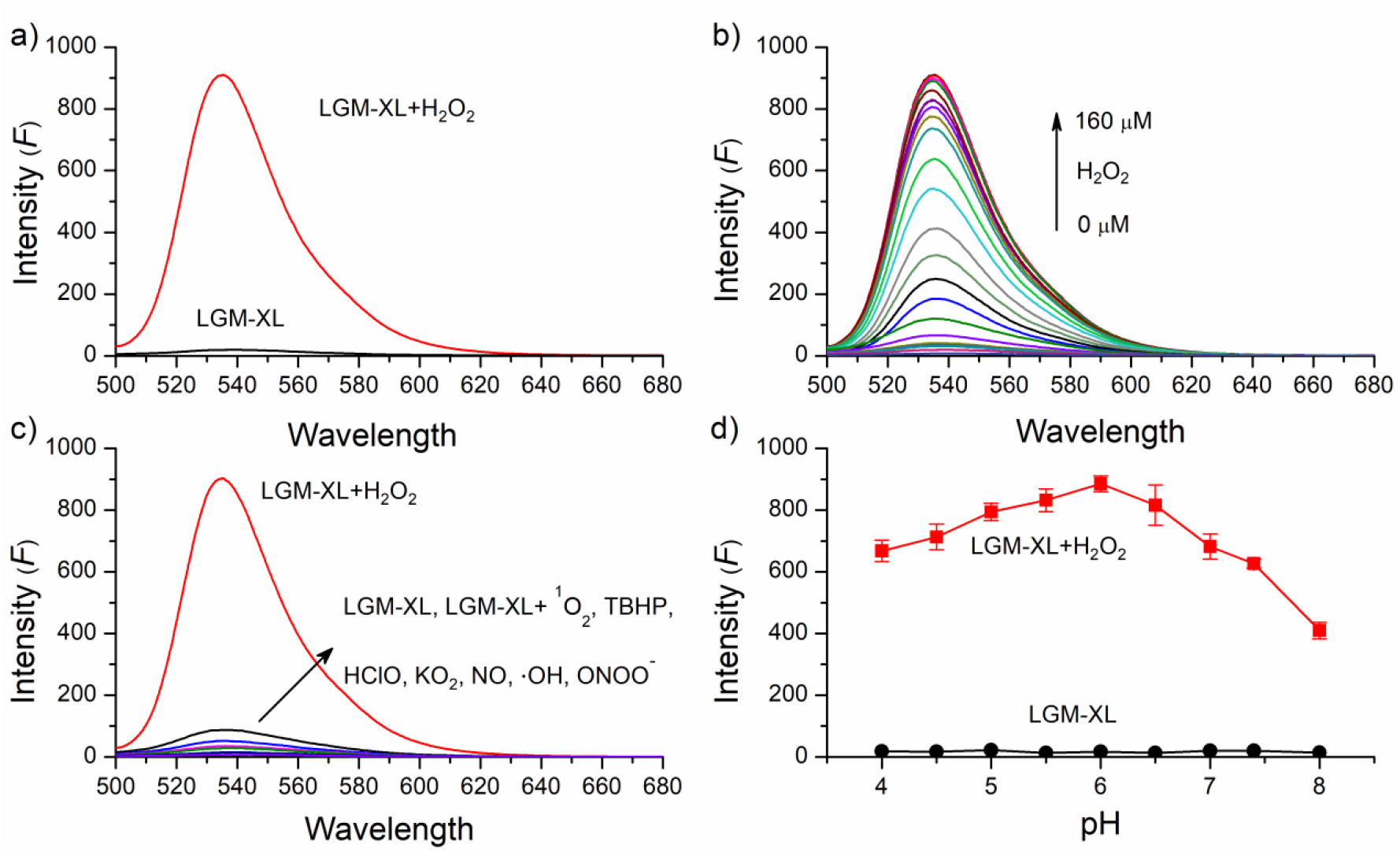
a) Fluorescence emission spectra of LGM-XL (1 μM) with or without incubation of H_2_O_2_ (160 μM) at 37 °C for 4 h in PBS (pH = 6.0). Excitation wavelength: 480 nm. b) The fluorescence emission spectra of LGM-XL (1 μM) in the presence of different concentrations of H_2_O_2_ at pH 6.0 (PBS). λ_ex_=480 nm. c) Fluorescence intensity (λ_em_ = 535 nm) of 1 μM LGM-XL toward various ROS. d). Fluorescence spectra of 1 μM LGM-XL to pH in the absence and the presence of 60 μM H_2_O_2_ in the buffer solution.

To ensure the accuracy of results, we further evaluated the selectivity and anti-interference of probe LGM-XL toward H_2_O_2_ in the presence of various relevant interfering substances including ROS reactive nitrogen species (RNS), biothiols and representative biologically relevant species in buffer solution (20 mM, pH 7.4) under physiological conditions. As shown in Figure 1c, only H_2_O_2_ caused a dramatic fluorescence enhancement, while the fluorescence signal was almost steady upon addition of other biologically relevant species, including hypochlorite (ClO),·OH, and ONOO^−^. These expermental results demonstrate that LGM-XL could be a specific fluorescent probe for the detection of H_2_O_2_ compared to other relevant analytes under complex physiological and biological conditions.

Subsequently, the pH working range of probe LGM-XL in the absence and presence of H_2_O_2_ was explored by recording fluorescence intensity. As is shown in Figure 1d, the metabolite solely showed good stability in all the buffers at pH 4-8 primarily; additionally, upon H_2_O_2_ addition, the fluorescence emission intensity at 535 nm was an obvious increase within the pH range of 4.5–7.4, indicating good biocompatibility of probe LGM-XL in both acid and alkaline environment, further enhancing the biological applications of probe in several cellular divisions within a biological scale of pH values. The above data proved that probe LGM-XL has a good capability for quantitative detection of H_2_O_2_ in vitro.

Next, the plateau of fluorescence intensity was collected after treatment of 9 equiv. of H_2_O_2_ under the excitation of 480 nm (Figure 2a). Additionally, the linear equation F = 15[LGM-XL] (μM) + 161 with a good linear relationship (R^2^=0.996) when the concentration of LGM-XL was increased from 0.2 μM to 1.8 μM. The detection limit (LOD) was then calculated as 0.21 μM based on a signal to noise ratio (S/N= 3), highlighting the superior sensitivity of probe LGM-XL for H_2_O_2_ detection among the reported data. Accordingly, the pseudo-first-order rate constant was determined to be 0.127 M^−1^ min^−1^. Additionally, a kinetic analysis of the LGM-XL reaction with H_2_O_2_ was also performed and the results were presented in Figure 2b. Remarkably, the fluorescence intensity ratio gradually increased and reached the balance point within 8 min after adding 160 μM H_2_O_2_ under the excitation of 535 nm, which indicated that the probe is advantageous for the real-time detection of H_2_O_2_ in the living body.

**Figure 2.**
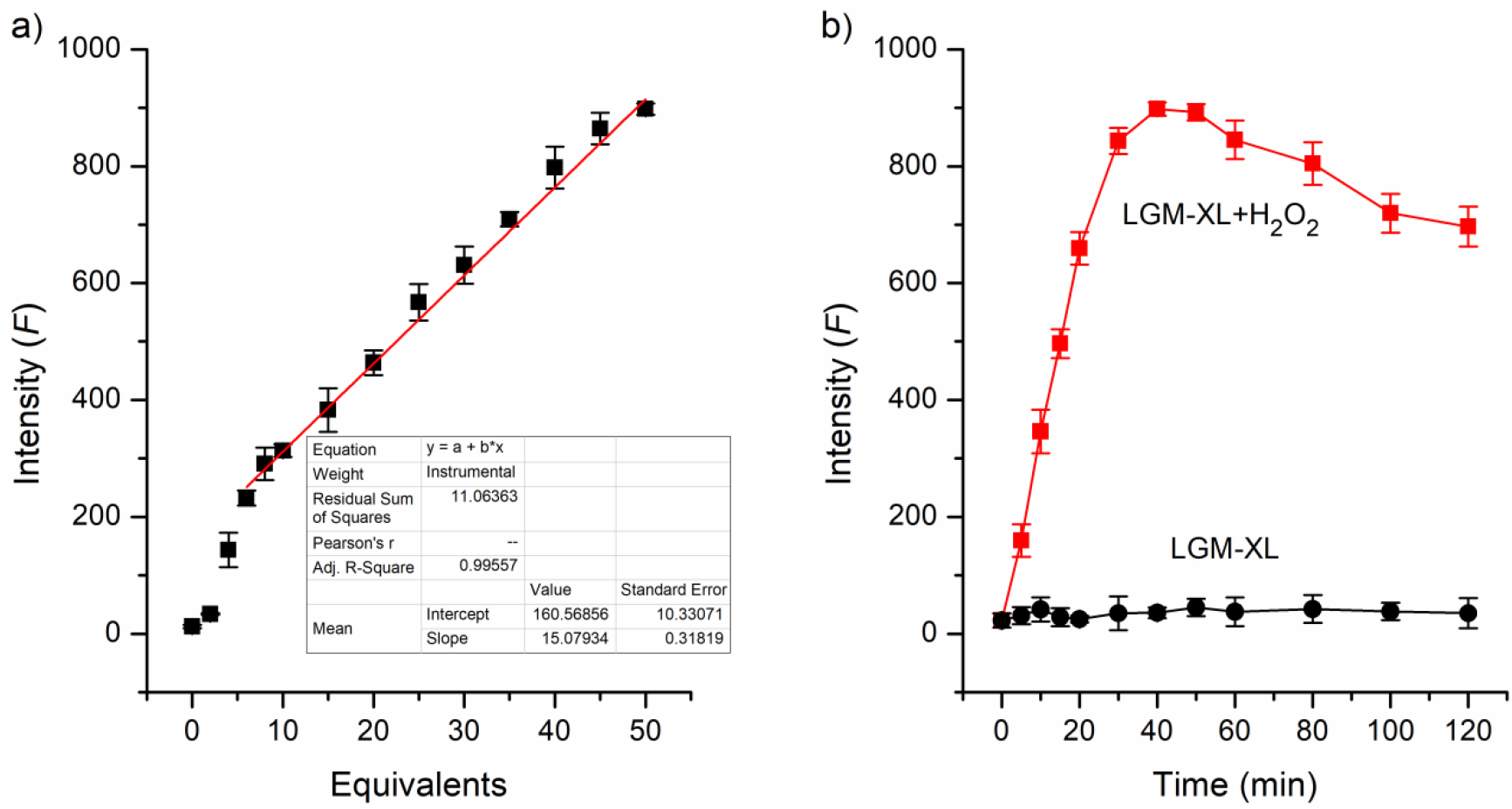
a) Plots of fluorescent intensity of LGM-XL (1 μM) as a function of H_2_O_2_ concentrations. b) Fluorescence intensity changes of LGM-XL (1 μM) exposed at pH 6.0 for different time periods. λ_ex_=480 nm.

### 3.2 The reaction mechanism of probe LGM-XL toward H_2_O_2_

For further validate the suggested mode of action of probe LGM-XL, the plausible recognizing mechanism of LGM-XLtoward H_2_O_2_ was proposed under the standard condition as described above (Scheme 2). When the probe LGM-XL interacts with H_2_O_2_, formed an unstable cyclic intermediate and released the fluorophore LGM. This hydrolysis-triggered substitution-cyclisation-elimination reaction recover the intramolecular charge process of the product, which can make the probe fluoresce at 535 nm. After the cyclization reaction and release of 5′-DFUR moiety, and as a result, released the 4-hydroxy-1,8-naphthalimide fluorophore. Whereupon the reaction product was observed and obtained by the spectra of the fluorescence of LGM-XL itself and fluorophore, indicating that probe reacts with H_2_O_2_ to produce the product of cyclization LGM. Additionally, a mixture of probe (1 μM) and H_2_O_2_ (10 μM) in PBS solutions was confirmed by the mass spectral analysis (Figure S5). Upon the addition of H_2_O_2_, a distinct peak at m/z = 994.3 (calcd. 993.26 for [M]^+^) as well as the naphthalimide byproduct ([M + H]^+^, m/z 399.1) were found, which strongly implied that the oxidative cleavage of LGM-XL by H_2_O_2_. Therefore, all these data are

**Scheme 2.**
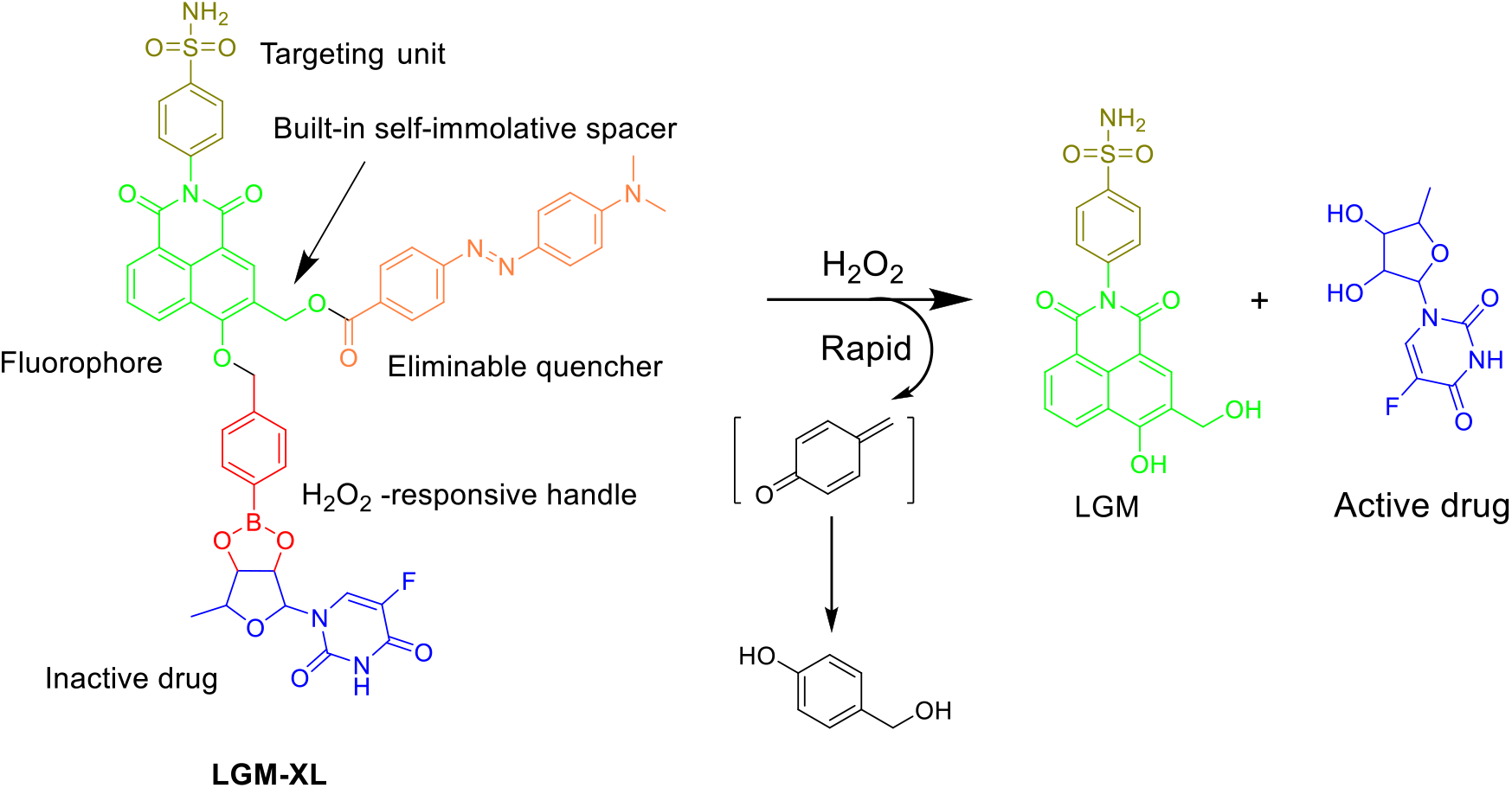
The structure and fluorescence response mechanism of LGM-XL.

### 3.3 Golgi-targeting capability of LGM-XL

Considering the excellent detection performance of LGM-XL, the connection between cell inflammation and the expression of the H_2_O_2_ were further investigated in several types of living tumor cells (HepG2, and RAW264.7 cells). we employed commercial e commercial probe Golgi-Tracker Red to co-stain with the probe (λ_ex_= 480 nm, λ_em_ = 535 nm) under confocal laser scanning microscopy (CLSM). As depicted in Figure 3, we treated the tumor cells with LGM-XL (1 µM) in culture medium at 37 °C for 10 min and then colocalization imaging. The results of confocal microscopy imaging showed that the fluorescent signals of probe could colocalize strongly with the fluorescence of Golgi-Tracker Red in Golgi, and the colocalization of Pearson coefficient was as high as 0.98 (HepG2 cells), and that from RAW264.7 cells were slightly lower (0.95). The difference in emission among these cell lines demonstrate that the selective uptake of LGM-XL by HepG2 cells through receptor-mediated endocytosis. Therefore, co–localization study proved that probe LGM-XL had the superior Golgi apparatus -targeting ability and the strong affinity during various pathological processes.

**Figure 3.**
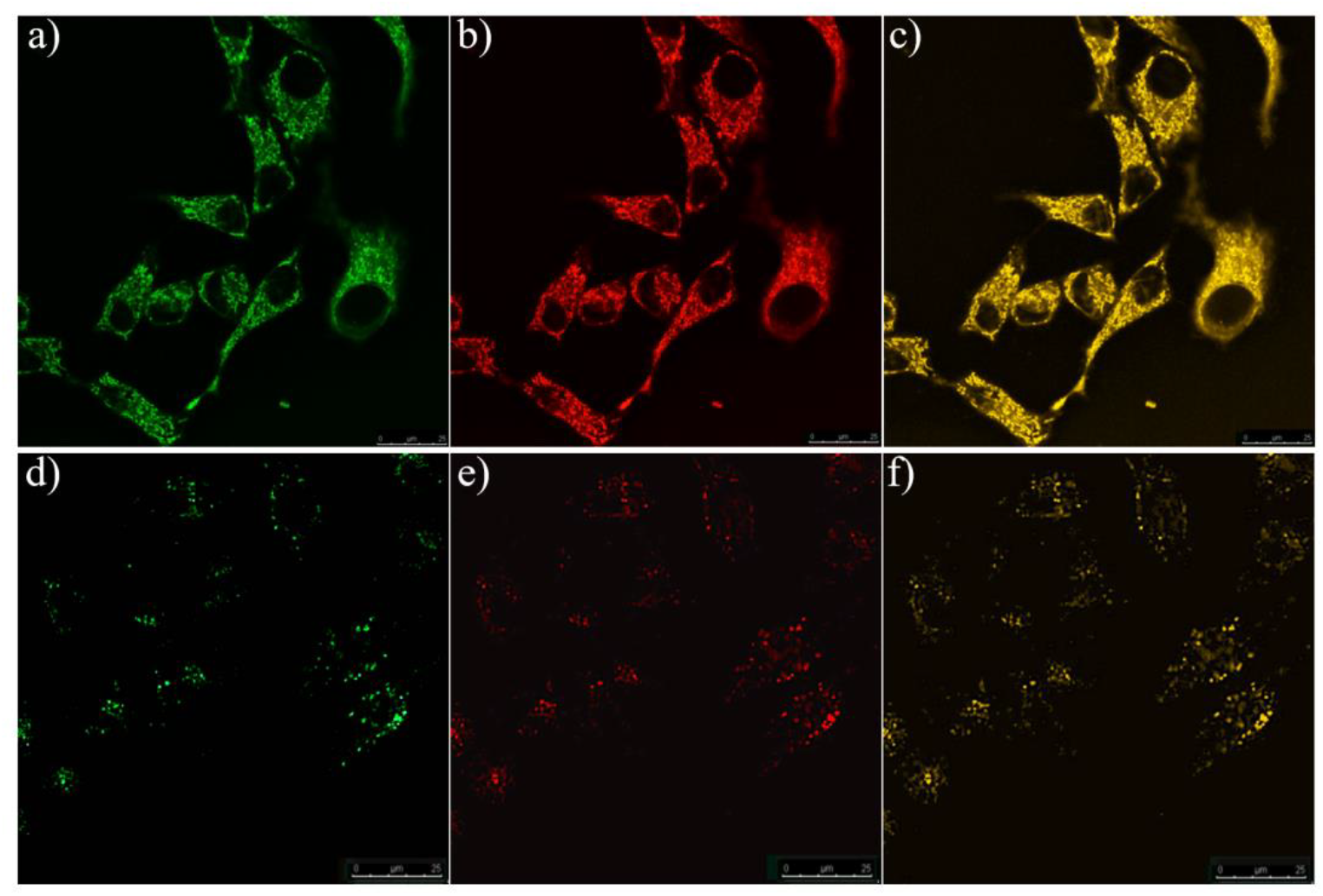
Co-localization cell imaging of LGM-XL (1 μM) and commercial dyes in HepG2 cells and RAW264.7 cells.

**Figure 4.**
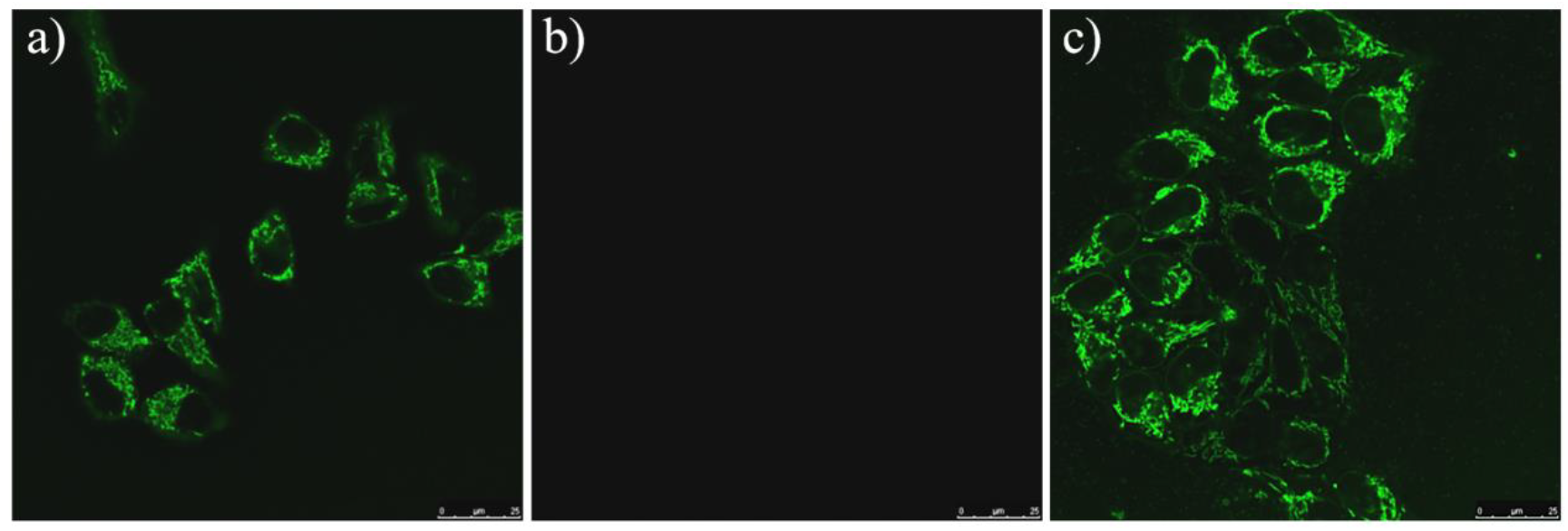
Co-localization cell imaging of LGM-XL (1 μM) and commercial dyes in HepG2 cells and RAW264.7 cells.

### 3.4 The recognition of probe LGM-XL to H_2_O_2_ in cellula

Next, to explore the relationship between H_2_O_2_ and Golgi stress response, we evaluated the total abundance and signal variations of LGM-XL on the change of H_2_O_2_ concentration at the Golgi. Given that the interference from cytoplasmic H_2_O_2_ during the Golgi stress response in living cells, LGM-XL was exposed to the intracellular H_2_O_2_ scavengers NAC (N-acetylcysteine) at first, and then the cells were tracked with different levels of Mone for 16 h. Monensin (Mone), a Golgi stress-inducing agent, was reported as the specific expression vector for endogenous H_2_O_2_ within the Golgi apparatus.^[30]^ When pretreating HepG2 cells with 0.5 mM NAC (H_2_O_2_ eliminator) for 30 min, the cells showed that no noticeable fluorescence signal in green channel was observed. By contrast, after the treatment of Monensin (10 μM for 6 h), the cells exhibited an enhanced fluorescence compared to the control group, indicating that the probe can be employed to detect the H_2_O_2_ activity in Golgi oxidative stress.

Figure 5 demonstrated that the encapsulation efficiency change curve of 5’-deoxy-5-fluorouridine (5′-DFUR) depended on the amount of H_2_O_2_ in PBS solution. Simulating the acid-triggered degradation of microenvironment, LGM-XL exhibited the highest release rate of 86% within 30 min at tumor pH 6.2 compared with that at physiological pH 7.4. These prove to be a beneficial feature that can improve the stability and biocompatibility in blood circulation using LGM-XL encapsulation in tumor therapy. Prior to cell imaging experiment, we first carried out standard MTT assays to ascertain the cytotoxicity test of probe LGM-XL in living HepG2 cells. As shown in Figure 5b, The HepG2 cells were incubated with LGM-XL at a lower concentration gradient (0, 0.2, 0.5, 1, 2, 5, 10, 20 and 30 µM) for 24 h. To our delight, this result demonstrated that the cell survival rates remained more than 85% even the concentration of the probe was up to 30 M after incubation for 24 h, indicating low cytotoxicity and good biocompatibility of this probe to living cells.

**Figure 5.**
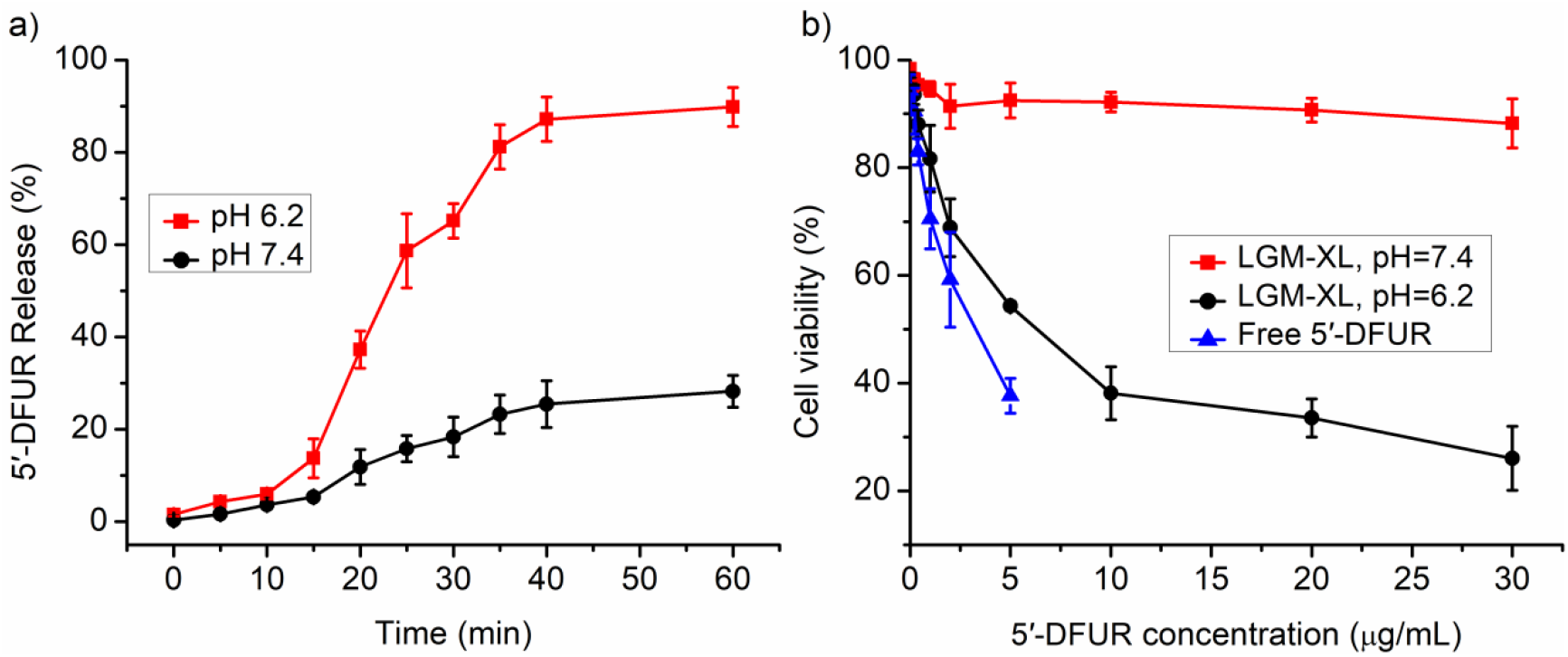
(a) In vitro release profiles of 5′ -DFUR from prodrug in PBS under different pH conditions. (b) In vitro cell viability assay of LGM-XL (1 μM) in HepG2 cells at pH 7.4 and 6.2 for 48 h.

### 3.5 Golgi oxidative stress and hypertension

Subsequently, the mouse inflammatory model was established by injection of lipopolysaccharide (LPS) in this study (Figure 6). Comparing the normal area, a large fluorescence turn-on signal was displayed at the area of acute inflammation after 20 min in the LPS-pretreated group. Thus, the above results revealed that probe LGM-XL could detect and image endogenous H_2_O_2_ in inflamed mice in situ, which may provide a potential method for the diagnosis of inflamed-related diseases.

**Figure 6.**
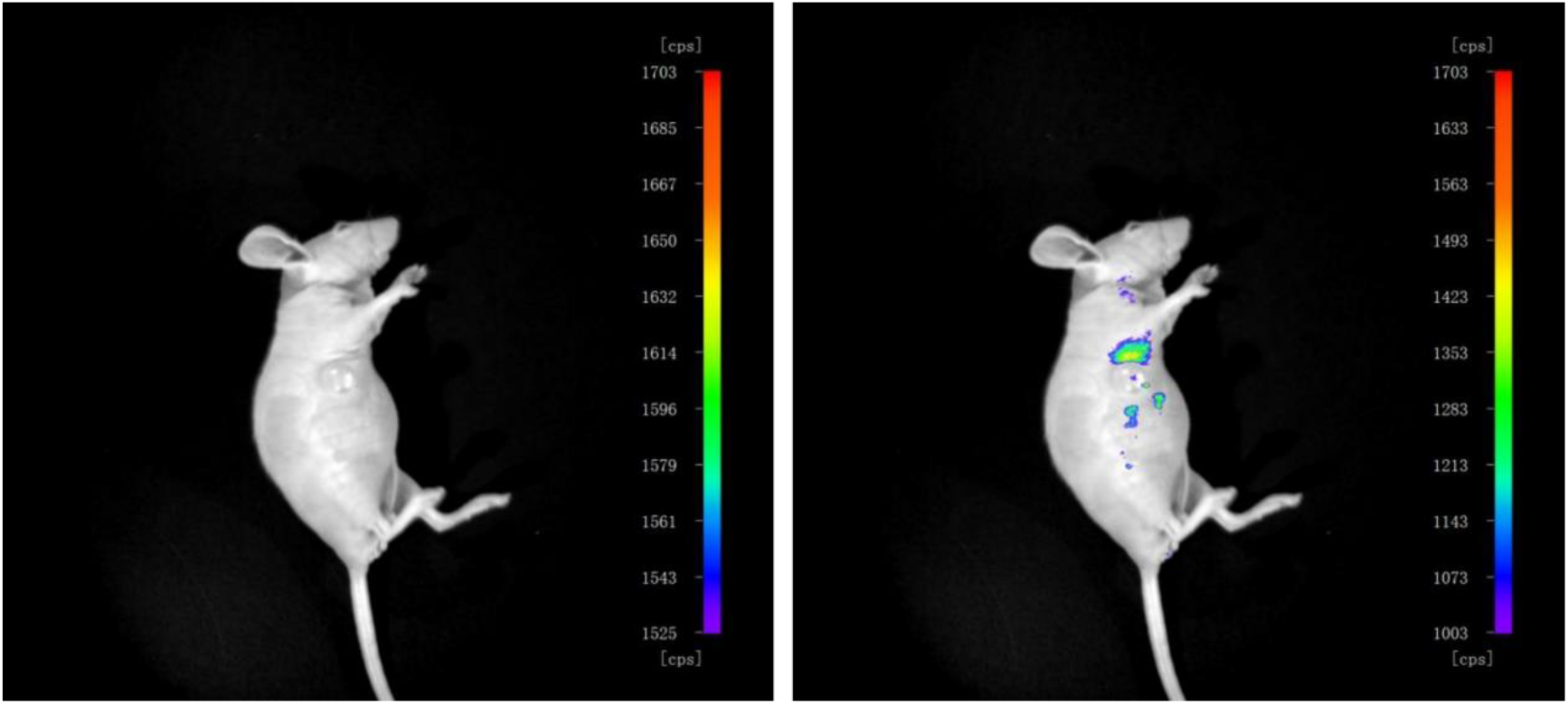
(a) In vivo imaging of tumor tissues in HepG2-cell-inoculated xenograft mice. The mice were tail-vein injected with 100 mM LGM-XL and lipopolysaccharide (LPS) in 200 mL of PBS. The fluorescent image of xenograft mice were taken after 3 injections.

Driven by the promising performance of LGM-XL in cells imaging, we wondered whether the probe bioimage endogenous H_2_O_2_ during the process of Golgi stress response in inflammatory mice. BALB/c nude mice used in vivo imaging experiments model were provided by commercial suppliers. The mice were subcutaneously injected with saline (20 μM) and probe LGM-XL (20 μM), respectively. The images were conducted after the incubation time of 10 min. As shown in Supplementary Figure S6, the mice exhibited significantly lower blood pressure (mSP (64.44 mmHg), mDP (49.13 mmHg), mAP (54.24 mmHg) compared with the control mice (mSP (77.62 mmHg), mDP (65.62 mmHg), mAP (69.62 mmHg)). These results validated the mice with hypertension caused severe Golgi oxidative stress by stimulation of LGM-XL.

Our attention then turned to the changes of body weight and tumor volume in hypertensive rats under three treatment regimens: (i) saline; (ii) 5′-DFUR; and (iii) a combination treatment of 5′-DFUR and Navitoclax (5′-DFUR + Nav). Nav is a senolytic agent to selectively eliminate senescent cells and inhibit tumor growth.^[31]^ As a control, saline-treated mice showed a slight change in mice body weight after 20 days treatment (Figure 7a). While the drug treated groups, the tumor weight was found significantly inhibited by the combination of 5′-DFUR+Nav. Meanwhile, Figure 7b showed that continuous injection of 5′-DFUR+Nav effectively reduced its volume, which was much better than saline or pure 5′-DFUR-treated mice. Overall, these results validated the AND-gate strategy could improve tumor accumulation effect during fluorescence-guided surgery by stimulation of LGM-XL.

**Figure 7.**
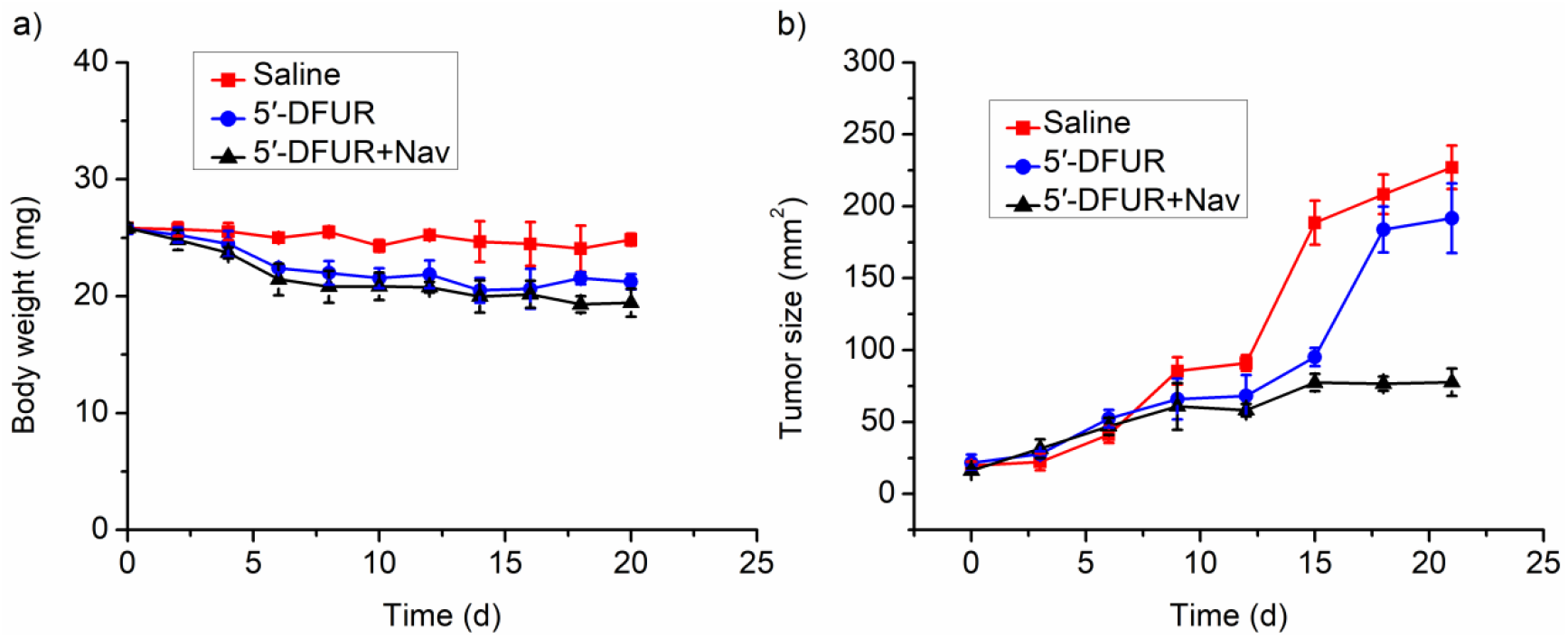
(a) Body weight curves in 20 days after intravenous injection with saline (control), 5′-DFUR and 5′-DFUR+Nav. (b) Tumor growth curves in 20 days after intravenous injection with saline (control), 5′-DFUR and 5′-DFUR+Nav.

## 4. Conclusions

To summarize, this work provided a robust tool for uncovering the relationship between Golgi oxidative stress and hypertension, as well as offers a method for unique interactions with LGM-XL with oxidative stress activation in early cancer cell diagnosis. The targeting system is based on the modification of naphthalimide fluorophore with ultra-high specificity and fast responses for Golgi H_2_O_2_ in cancer cells during oxidative stress. By utilization of LGM-XL, we successfully modulated H_2_O_2_ levels in the oxidative stress-induced kidneys injury mouse model with hypertension. Additionally, the present strategy showed a significantly prolonged circulation time that would reduce the side effects of the drug. These compelling results underscore the promising potential of LGM-XL for designing oxidative stress-responsive prodrug of novel therapeutics to treat H_2_O_2_-related diseases (e.g., Hypertension) in the future.

## Acknowledgment

We would like to acknowledge the financial support of the National Natural Science Foundation of China (NSFC) projects (22267007), the Longyuan Youth Innovation and Entrepreneurship Talent Project of Gansu Province (2023LQGR15).

